# GABAergic network from AVP neurons to VIP neurons in the suprachiasmatic nucleus sets the activity/rest time of the circadian behavior rhythm

**DOI:** 10.1101/2025.04.28.650944

**Authors:** Yubo Peng, Yusuke Tsuno, Takashi Maejima, Mohan Wang, Ayako Matsui, Michihiro Mieda

**Affiliations:** Department of Integrative Neurophysiology, Graduate School of Medical Sciences, Kanazawa University, Kanazawa, Japan

**Keywords:** circadian rhythm, biological clock, suprachiasmatic nucleus, GABA, CRISPR-Cas9, fiber photometry, optogenetics

## Abstract

The central circadian clock of the suprachiasmatic nucleus (SCN) is a network composed of multiple types of γ-aminobutyric acid (GABA)-ergic neurons and glial cells. However, the precise role of GABAergic transmission in the SCN remains unclear. In this study, we investigated the GABAergic regulation from arginine vasopressin (AVP)-producing neurons in the SCN shell to vasoactive intestinal polypeptide (VIP)-producing neurons in the SCN core. Blocking GABA release from AVP neurons by a vesicular GABA transporter (*Vgat*) gene deletion lengthened the activity time (the interval between the onset and offset of locomotor activity) and shortened the duration of high Ca^2+^ activity in VIP neurons to match the behavioral rest time. Conversely, eliminating functional GABA_A_ receptors (GABA_A_R) in VIP neurons by in vivo genome editing reduced locomotor activity level and the activity time, and lengthened the high Ca^2+^ duration in VIP neurons. Optogenetic activation of AVP neurons in vivo increased Ca^2+^ in VIP neurons during the night. A similar Ca^2+^ response of VIP neurons to AVP neuronal activation was also observed in SCN slices and was inhibited by a GABA_A_R antagonist, gabazine. Importantly, gabazine application alone raised the baseline Ca^2+^ in VIP neurons, suggesting a tonic depression of these neurons by GABA. Moreover, AVP neuronal activation decreased Ca^2+^ in non-AVP neurons located between AVP- and VIP-rich regions in the SCN. These results suggest that GABA from AVP neurons disinhibits VIP neurons indirectly by suppressing other intermediate GABA neurons to set the behavior activity/rest time precisely.

## Introduction

The suprachiasmatic nucleus (SCN) of the hypothalamus serves as the central circadian clock in mammals, orchestrating multiple circadian biological rhythms in the body [1]. The SCN is composed of approximately 20,000 cells. Individual SCN cells can generate cellular circadian oscillations driven by the autoregulatory transcription-translation feedback loop (TTFL) of clock genes, including *Bmal1*, *Clock*, *Per1/2/3*, and *Cry1/2*. Interestingly, it is not only the SCN cells that share TTFL-driven cellular clocks but also most cells throughout the body. Rather, intercellular communication between SCN cells is essential to generate a highly robust, coherent circadian rhythm [1,2].

The SCN is a network of neurochemically heterogeneous γ-aminobutyric acid (GABA)-ergic neurons. Several types of GABA neurons can be distinguished by the co-expressed neuropeptides [1,2]. Arginine vasopressin (AVP)-producing GABAergic neurons in the SCN shell, the dorsomedial part of the SCN, and vasoactive intestinal polypeptide (VIP)-producing GABAergic neurons in the SCN core, the ventrolateral part, are the two of the major neuron types. VIP has been reported as a critical factor in the maintenance and synchronization of SCN neurons [3–6]. In addition, these neurons also contribute to the output pathway from the SCN [7–9]. In contrast, AVP neurons have been suggested to be the primary pacesetter cells that determine the period length of the circadian rhythm generated by the SCN network [10–13].

The functional roles of GABA-mediated signaling in the SCN network remain controversial [14]. GABA was initially reported to synchronize action potential firing rhythms of dispersed SCN neurons through GABA_A_ receptors (GABA_A_R) [15]. Studies using SCN slices and GABA_A_R antagonists showed that GABA synchronizes or desynchronizes the cellular circadian oscillations depending on the lighting conditions before the slice preparation [16–18]. In addition, the regulation of extracellular GABA in the SCN by astrocytes was reported to be critical for circadian timekeeping in neonatal SCN cultures [19–21]. The SCN-specific in vivo deletion of the vesicular GABA transporter gene (*Vgat*, also called *Slc32a1*), which is necessary for filling synaptic vesicles with GABA and thus for synaptic GABA release [22], attenuated the circadian behavior rhythm but remained the TTFL oscillation more or less normal [23,24].

We previously demonstrated that AVP neuron-specific deletion of *Vgat* drastically alters the daily locomotor activity pattern without changing the free-running period [25]. Namely, these mice showed a marked lengthening of the activity time in the circadian behavior rhythm due to an extended interval between morning and evening locomotor activities. GABA released by AVP neurons was suggested not to significantly affect the synchrony of TTFLs among SCN neurons but to regulate the phase relationships between TTFLs in the SCN and circadian morning/evening locomotor activities [25]. In addition, the daily rhythm of the in vivo multiunit activity (MUA) in the SCN clearly changed to an aberrant bimodal pattern that correlated with dissociated morning/evening locomotor activities [25]. These observations indicated that GABAergic transmission from AVP neurons regulates the activities of other SCN neurons to temporally restrict circadian behavior to appropriate time windows in SCN TTFLs [25]. In this study, we show the critical role of indirect GABAergic regulation of AVP neurons on VIP neurons in the activity/rest time setting.

## Results

### In vivo Ca^2+^ rhythm of SCN VIP neurons tracks the compressed rest period of the behavior rhythm in AVP neuron-specific *Vgat* deficiency

AVP neuron-specific *Vgat*-deficient mice (*Avp-Vgat^−/−^*) showed a lengthened activity time (the interval between the locomotor activity onset and offset) of behavior rhythm with little change in the intracellular Ca^2+^ rhythm of AVP neurons, resulting in a misalignment of locomotor activity and AVP neuronal Ca^2+^ rhythm [25]. To elucidate how GABA released from AVP neurons regulates the activity of VIP neurons in the SCN, we first examined the temporal relationship between the circadian behavioral rhythm and the intracellular Ca^2+^ in VIP neurons (VIP-Ca^2+^) of *Avp-Vgat^−/−^* mice in vivo using fiber photometry.

To specifically target the jGCaMP7s expression in VIP neurons of *Avp-Vgat^−/−^* mice, we introduced a *Vip-tTA* allele [26] into *Avp-Cre; Vgat ^flox/flox^* (*Avp-Vgat^−/−^*) or *Avp-Cre; Vgat ^wt/flox^* (control) mice. In *Vip-tTA* mice, the tetracycline transactivator (tTA) is expressed specifically in VIP neurons. We then specifically expressed the fluorescent Ca^2+^indicator jGCaMP7s [27] in SCN VIP neurons by focally injecting a tTA-dependent AAV vector (AAV-*TRE-jGCaMP7s*) and implanted an optical fiber just above the SCN (Fig 1A and 1B) [12].

**Figure 1.**
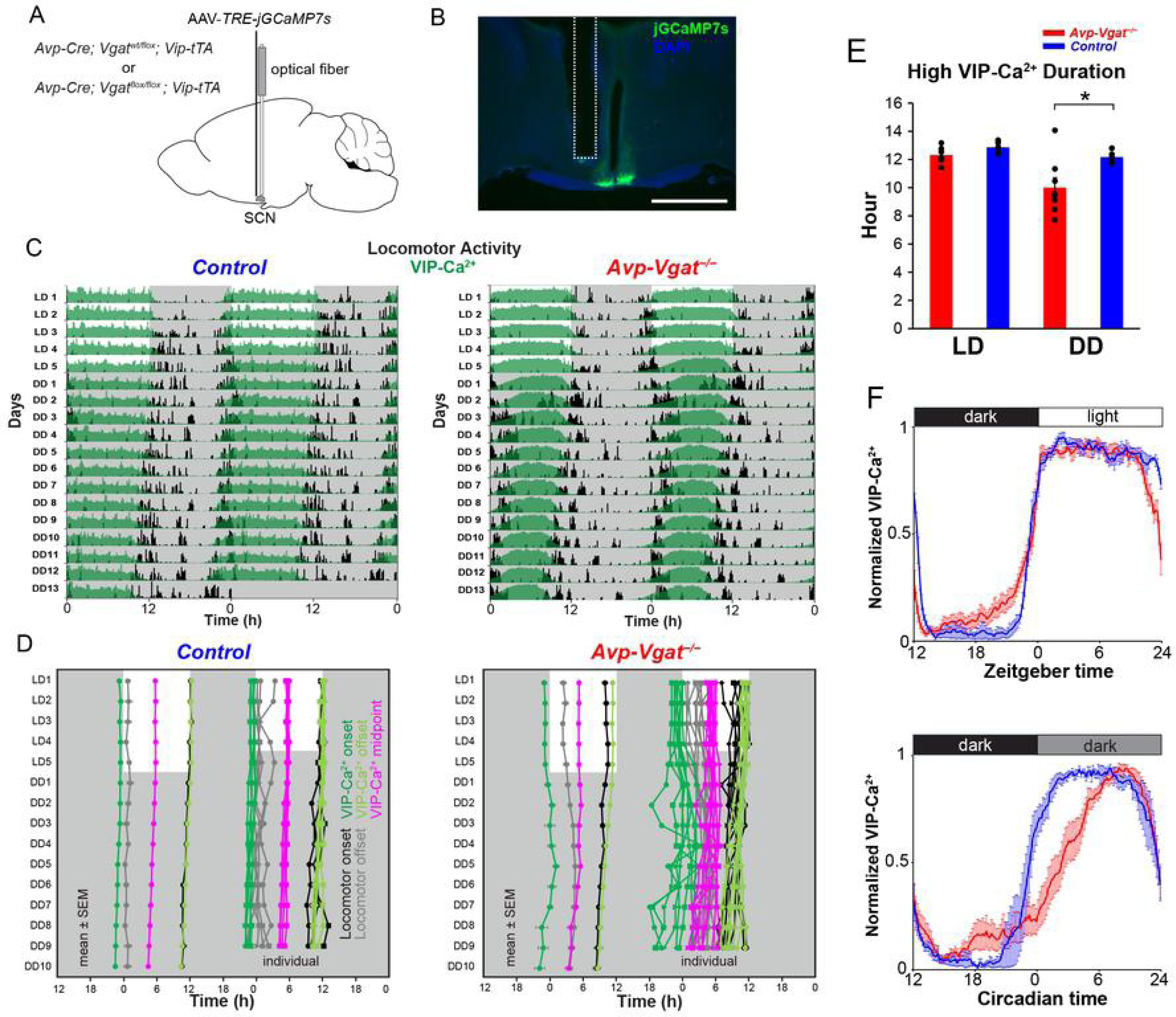
VIP-Ca^2+^ rhythm matches the compressed rest time of the locomotor activity in *Avp-Vgat^−/−^*mice. (A) Schematic diagram of viral vector (*AAV-TRE-jGCaMP7s*) injection and optical fiber implantation at the SCN in control (*Avp-Cre; Vgat ^wt/flox^; Vip-tTA*) or *Avp-Vgat^−/−^* (*Avp-Cre; Vgat ^flox/flox^; Vip-tTA*) mice for fiber photometry recording. (B) A representative coronal section of mice with jGCaMP7s expression in SCN VIP neurons. A white dotted square shows the estimated position of the implanted optical fiber. Green, jGCaMP7s; blue, DAPI. Scale bar, 1 mm. (C) Representative plots of the in vivo jGCaMP7s signal of SCN VIP neurons (green: VIP-Ca^2+^) overlaid with locomotor activity (black) in actograms. Control (Left) and *Avp-Vgat^−/−^* (Right) mice were initially housed in LD (LD1 to LD5), then in DD (DD1 to DD13). Gray shading indicates the time when the lights were off. (D) Plots of locomotor activity onset (black), activity offset (gray), VIP-Ca^2+^ onset (green), VIP-Ca^2+^ offset (light green), and VIP-Ca^2+^ midpoint (magenta) of mean ± SEM (left column) and individual (right column) in control and *Avp-Vgat^−/−^* mice. Identical marker shapes indicate data from the same animal. (E) High VIP-Ca^2+^ duration in LD (LD1-5, left) or in DD (DD6-10, right). (F) Normalized VIP-Ca^2+^ daily rhythm profiles in LD (LD1-5, top) or in DD (DD6-10, bottom). In DD, CT12 was determined as the onset of locomotor activity. Red, *Avp-Vgat^−/−^* (n = 8); blue, Control (*Avp-Vgat^+/−^*, i.e., *Avp-Cre; Vgat ^wt/flox^*, n = 5); Values are mean ± SEM. *P < 0.05 by two-tailed Welch’s t-test.

The VIP-Ca^2+^ rhythm was synchronized antiphasically with the locomotor activity rhythm in control (*Avp-Cre; Vgat^wt/flox^; Vip-tTA*) mice (Fig 1C and 1D) and showed high levels of Ca^2+^ across the entire span of the behavioral rest period, as reported previously [12,28–30]. In *Avp-Vgat^−/−^* mice, in contrast, the duration of high VIP-Ca^2+^ was clearly compressed (*Avp-Vgat^−/−^*, 10.0 ± 0.7 h; Control, 12.2 ± 0.2 h, P = 0.02) complementarily to the expansion of behavioral activity time (*Avp-Vgat^−/−^*, 19.2 ± 0.3 h; Control, 13.6 ± 0.4 h, P < 0.001) in constant darkness (DD) condition (Figs 1C-1F and S1). The VIP-Ca^2+^ offset almost coincided with the locomotor activity onset (Fig 1D).

These results suggest that the temporal relationship between the VIP-Ca^2+^ rhythm and the locomotor activity is essentially maintained in the *Vgat*-deficiency of AVP neurons and that GABA from AVP neurons regulates the VIP-Ca^2+^ rhythm.

### Elimination of GABA_A_R in SCN VIP neurons reduces locomotor activity and shortens the activity time of the behavior rhythm

Thus, SCN VIP neurons likely receive direct or indirect GABAergic regulation from AVP neurons, as well as GABA from other types of SCN neurons. To investigate the role of GABAergic signaling in VIP neurons in generating the behavior rhythm, we next aimed to eliminate ionotropic GABA_A_R specifically in these neurons. GABA_A_R is essentially a pentameric protein composed of α, β, and γ or δ subunits [31]. Each subunit has multiple subtype genes, but we had no information about which genes to delete. Considering that the β subunit is necessary for functional GABA_A_Rs and that only three subtype genes encode the β subunit (*Gabrb1*-*3*), we introduced indel mutations in all three simultaneously and specifically in VIP neurons by in vivo genome editing. We first crossed Cre-dependent SpCas9-expressing (*Rosa-LSL-SpCas9-2A-EGFP*) mice [32] with VIP neuron-specific Cre driver (*Vip-ires-Cre*) mice [33]. Then we injected the mixture of three AAV vectors, each expressing gRNAs targeting one of the *Gabrb* genes (AAV-*U6-gGabrb1, 2, 3-EF1α-DIO-mCherry*), into the SCN of these mice (*Vip-GABA_A_R^−/−^* mice) (Figs 2A, 2B, S2A and S2B). Indeed, GABA_A_R-mediated postsynaptic currents (GPSCs) disappeared almost completely in VIP neurons of SCN slices prepared from *Vip-GABA_A_R^−/−^* mice, confirming the effectiveness of this method (S2C and S2D Figs).

**Figure 2.**
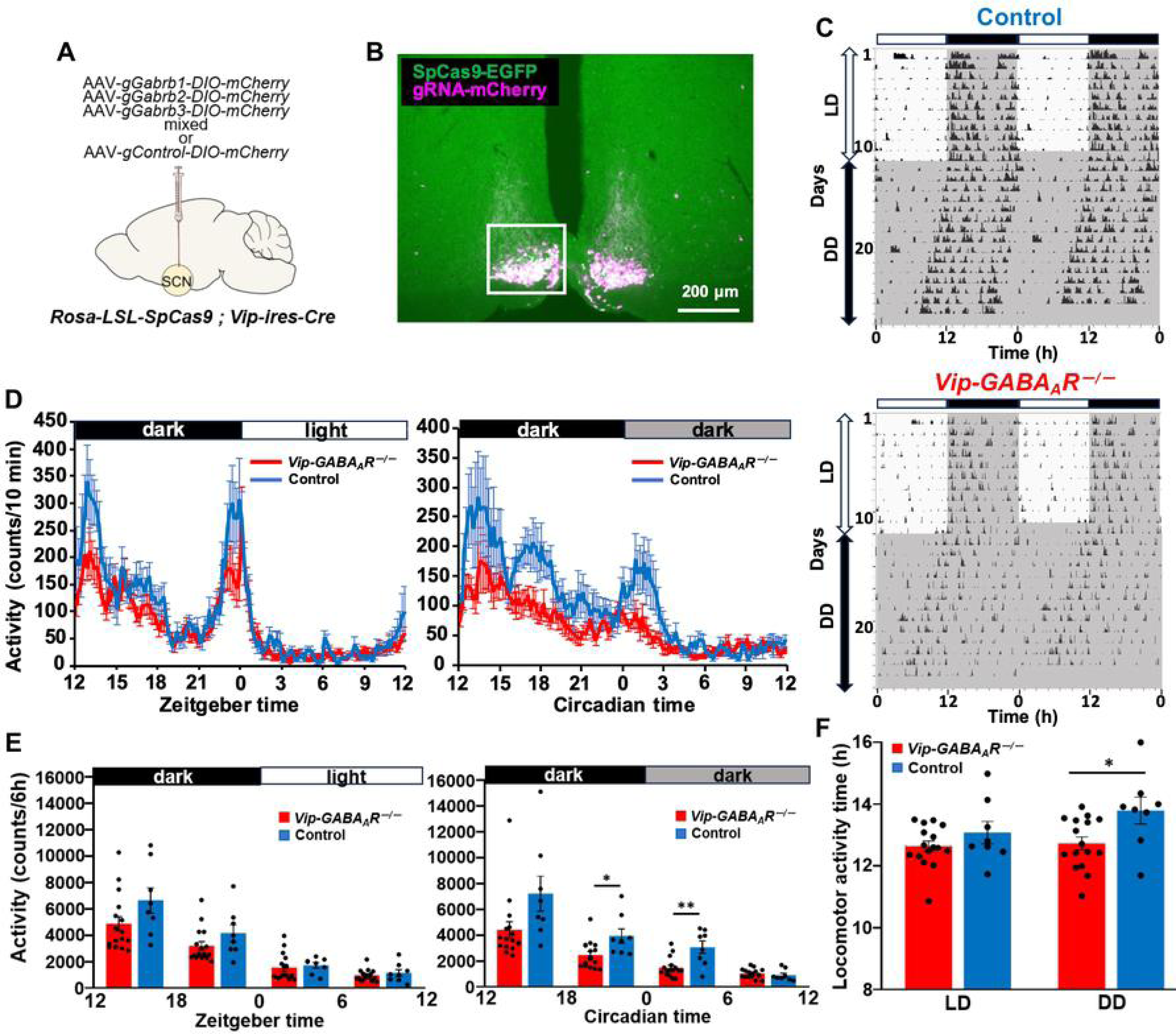
*Vip-GABA_A_R ^−/−^* mice reduce the locomotor activity and shorten the activity time. (A) Schematic diagram of viral vector (AAV-*U6-gGabrb1,2,3-EF1α-DIO-mCherry* or AAV-*U6-gControl-EF1α-DIO-mCherry*) injection at the SCN in *Rosa26-CAG-LSL-SpCas9-2A-EGFP; Vip-ires-Cre* mice to generate control or *Vip-GABA_A_R^−/−^* mice for locomotor activity recording. (B) A representative coronal section of mice with SpCas9-2A-EGFP and gRNA-DIO-mCherry expression in SCN VIP neurons. Green, SpCas9-EGFP; magenta, mCherry. (C) Representative locomotor activity of control and *Vip-GABA_A_R^−/−^*mice (home-cage activity). Gray shading indicates the time when the lights were off. (D) Averaged daily profile of locomotor activity in LD (left) or DD (right). (E) Average locomotor activity counts per 6-hour interval in LD (left) or DD (right). (F) Activity time of locomotor activity rhythm in LD (left) or in DD (right). Blue, Control; red, *Vip-GABA_A_R^−/−^*. Values are mean ± SEM. n = 8 for Control, n = 16 for *Vip-GABA_A_R^−/−^* mice. *P < 0.05; **P < 0.01 by two-way repeated measures ANOVA post-hoc two-tailed Student’s t-test with Bonferroni correction (E), or by two-tailed Student t tests (F).

In LD, *Vip-GABA_A_R^−/−^* mice showed a daily locomotor activity rhythm comparable to controls. In DD, however, the nocturnal locomotor activity of *Vip-GABA_A_R^−/−^* mice was significantly reduced with less obvious locomotor onset and offset without the bimodal morning and evening locomotor activities (Fig 2C-2E), resulting in a shortened activity time (*Vip-GABA_A_R^−/−^*, 12.73 ± 0.21 h; Control, 13.79 ± 0.43 h, P = 0.0189) (Fig 2F). In contrast, their free-running period and amplitude in DD did not alter significantly (S2E and S2F Figs). These results suggest that the disinhibition of VIP neurons due to the absence of GABA_A_Rs may suppress the locomotor activity in the subjective night.

### The high Ca^2+^ duration in VIP neurons is lengthened in *Vip-GABA_A_R****^−^****^/^****^−^*** mice

In *Avp-Vgat^−/−^* mice, the high VIP-Ca^2+^ duration of the VIP-Ca^2+^ rhythm was shortened while the behavioral activity time was lengthened. Therefore, we next investigated how the VIP-Ca^2+^ rhythm alters in vivo in *Vip-GABA_A_R^−/−^* mice (Fig 3A and 3B).

**Figure 3.**
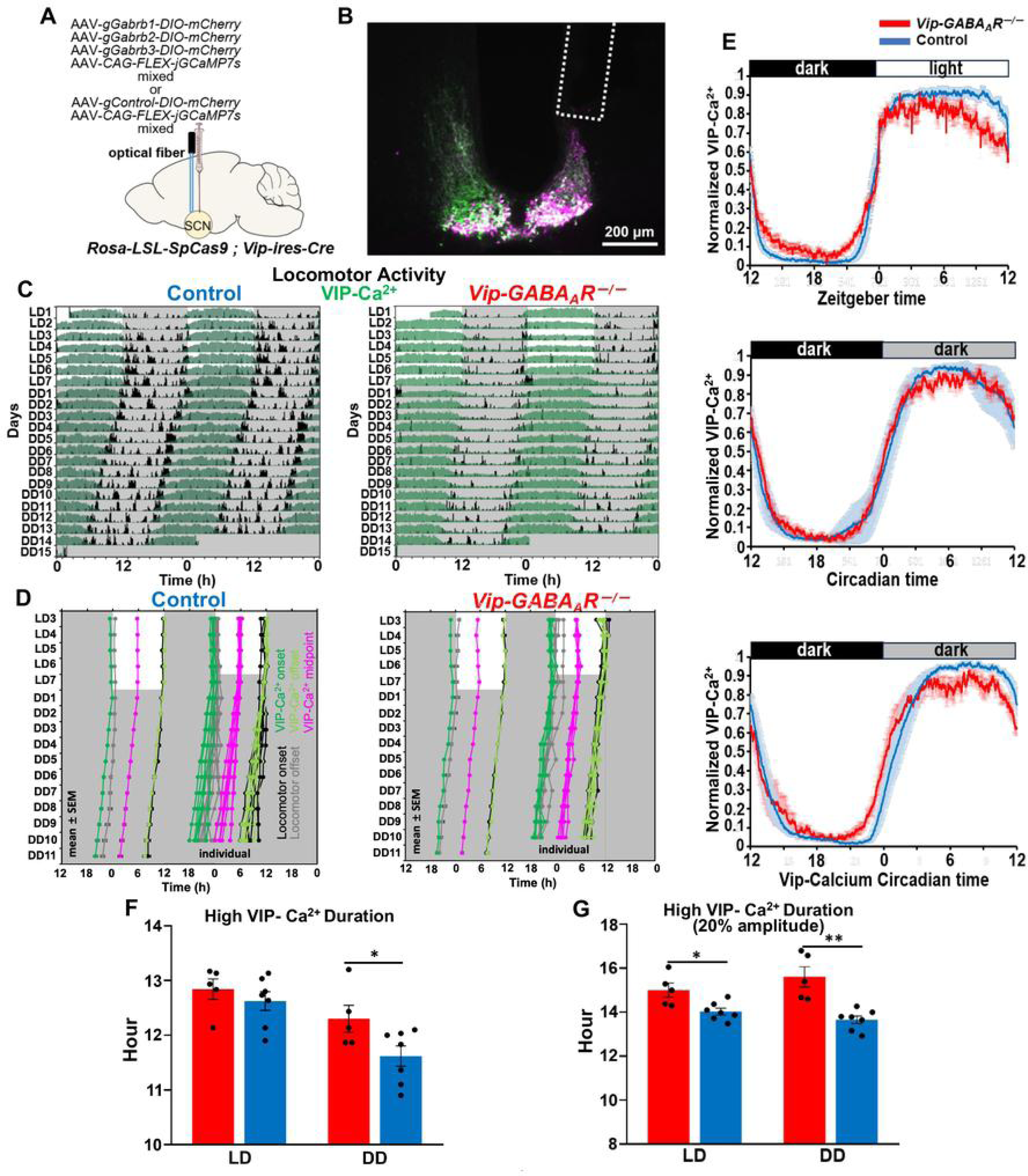
Duration of high VIP-Ca^2+^ lengthens in *Vip-GABA_A_R^−/−^* mice. (A) Schematic diagram of viral vector (AAV-*CAG-FLEX-jGCaMP7s* and AAV-*U6-gGabrb1,2,3-EF1α-DIO-mCherry* or AAV-*U6-gControl-EF1α-DIO-mCherry*) injection and optical fiber implantation at the SCN in *Rosa26-CAG-LSL-SpCas9-2A-EGFP; Vip-ires-Cre* mice to generate control or *Vip-GABA_A_R^−/−^* mice for fiber photometry recording. (B) A representative coronal section of mice with jGCaMP7s expression and gRNA-DIO-mCherry expression in SCN VIP neurons. A white dotted square shows the estimated position of the implanted optical fiber. Green, jGCaMP7s; magenta, mCherry. (C) Representative plots of the in vivo jGCaMP7s signal of SCN VIP neurons (green) overlaid with locomotor activity (home-cage activity) (black) in actograms. Control (Left) and *Vip-GABA_A_R^−/−^* (Right) mice were initially housed in LD (LD1 to LD7) and then in DD (DD1 to DD15). Gray shading indicates the time when the lights were off. (D) Plots of locomotor activity onset (black), activity offset (gray), VIP-Ca^2+^ onset (green), VIP-Ca^2+^ offset (light green), and VIP-Ca^2+^ midpoint (magenta) of mean ± SEM (left column) and individual (right column) in control and *Vip-GABA_A_R^-/-^* mice. (E) Normalized VIP-Ca^2+^ daily rhythm profiles in LD (LD3-7, top) or in DD (DD5-14,middle and bottom). In DD, circadian time 12 was determined as the onset of locomotor activity and VIP-Ca^2+^ circadian time 0 was determined as the onset of VIP-Ca^2+^. (F) High VIP-Ca^2+^ duration in LD (LD3-7, left) or in DD (DD5-14, right). (G) High VIP-Ca^2+^ duration in LD (LD3-7, left) or in DD (DD5-14, right) calculated with 20% amplitude of the VIP-Ca^2+^ profile as the threshold to determine the Ca^2+^ onset and offset. Values are mean ± SEM. n = 7 for control, n = 5 for *Vip-GABA_A_R^−/−^*mice. *P < 0.05; **P < 0.01 by two-tailed Student t tests.

In *Vip-GABA_A_R^−/−^* mice, the locomotor activity onset and offset became obscure due to the reduced activity levels (Fig 3C), as described earlier (Fig 2). Correspondingly, the high Ca^2+^ duration of the VIP-Ca^2+^ rhythm was significantly extended in these mice (*Vip-GABA_A_R^−/−^*, 12.30 ± 0.25 h; Control, 11.62 ± 0.19 h, P = 0.0483), which is consistent with the reduced behavioral activity time observed in these mice (Figs 3D-3F, S3B and S3C). The free-running period of the VIP-Ca^2+^ rhythm was comparable between genotypes (S3D Fig). Notably, the shape of the daily VIP-Ca^2+^ profiles appeared to be qualitatively different between *Vip-GABA_A_R^−/−^*and control mice (Fig 3E). In fact, the latter showed a relatively smooth quasi-rectangular wave, while the former fluctuated more on the plateau (Figs 3E and S3A). These observations might reflect the presumed higher basal Ca^2+^ level in VIP neurons lacking GABA_A_R, causing a ceiling effect on the plateau. Indeed, the high VIP-Ca^2+^ duration lengthening was more pronounced when the threshold for determining Ca^2+^ onset and offset was set at 20% of the amplitude (max-min) rather than 50%, as in other analyses (Fig 3E and 3G). These data suggest that the GABA_A_R elimination in VIP neurons causes a significant extension of high Ca^2+^ duration in vivo, which may subsequently suppress locomotor activity.

### Optogenetic activation of AVP neurons increases VIP-Ca^2+^ in a time-dependent manner in vivo

Previous studies have reported that AVP neuronal fibers make sparse contacts onto VIP neurons and that VIP neurons respond to the optogenetic activation of AVP neurons by increasing Ca^2+^ at around ZT22 in vivo [12,34]. Therefore, to further investigate the functional connectivity between AVP and VIP neurons, we next tested the time-of-day dependency of VIP-Ca^2^ response to the optogenetic activation of AVP neurons in vivo. To this end, ChrimsonR-mCherry, a red light-gated cation channel [35,36], and jGCaMP7s were expressed specifically in AVP and VIP neurons, respectively, by injecting AAV-*CAG-FLEX-ChrimsonR-mCherry* and AAV-*TRE-jGCaMP7s* into the SCN of *Avp-Cre; Vip-tTA* mice. Interestingly, optogenetic stimulation of AVP neurons increased VIP-Ca^2+^ only during the night, when basal Ca^2+^ is low, but not during the day, when basal Ca^2+^ is high (Fig 4A-4E). The most potent responses were observed at ZT14 and ZT22. These results suggest that AVP neurons can activate VIP neurons during the night in vivo.

**Figure 4.**
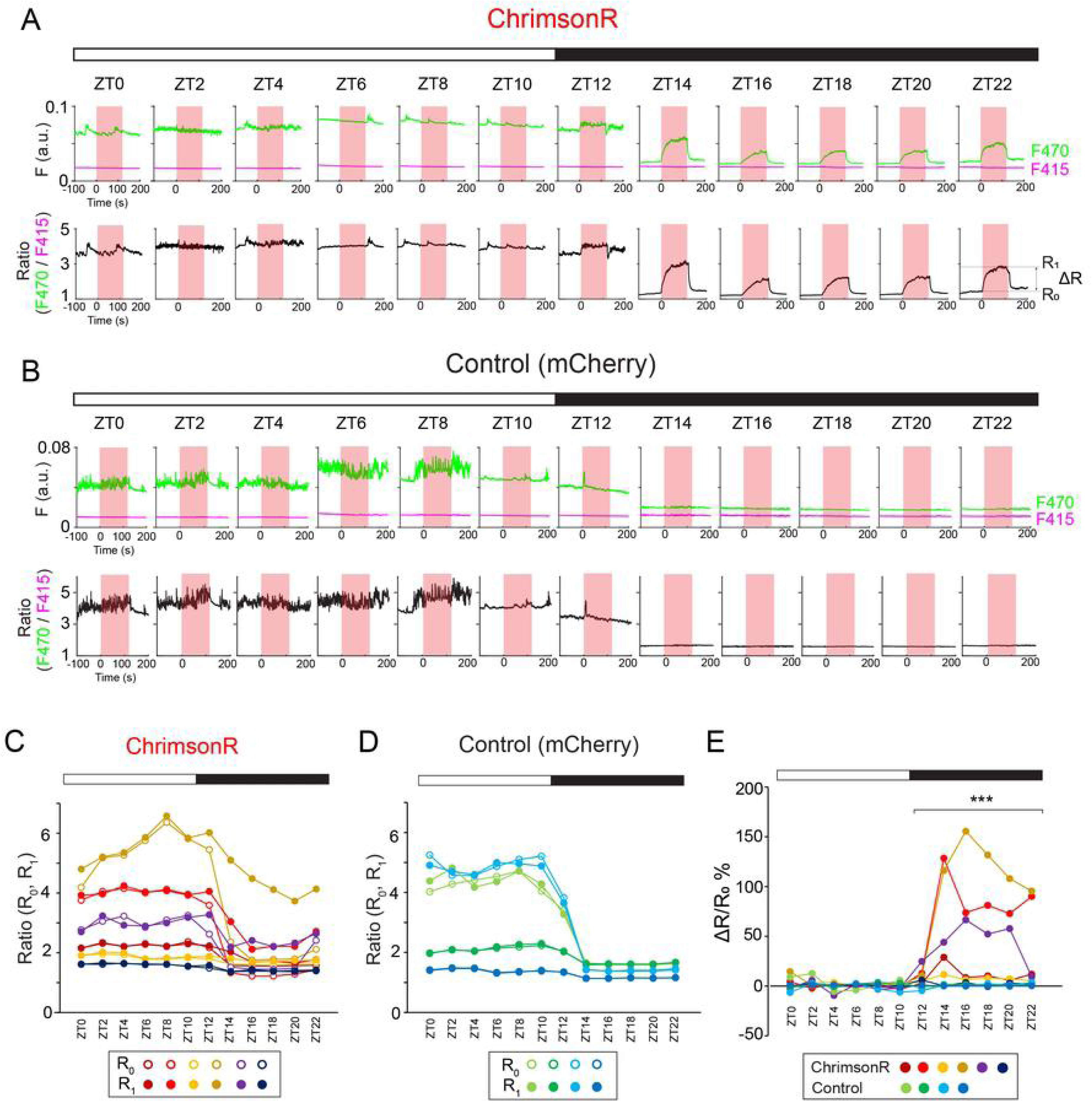
Optogenetic stimulation of AVP neurons increases VIP-Ca^2+^ in vivo during the night. (A, B) Top: Representative traces of the jGCaMP7s signal of SCN VIP neurons upon optogenetic stimulation of AVP neurons at various timing in vivo (A, ChrimsonR; B, mCherry-Control). Green traces indicate the fluorescence (F) value at the 470 nm light excitation (F470), Ca^2+^-dependent signal. Magenta traces indicate the fluorescence value at the 415 nm light excitation (F415), Ca^2+^-independent control signal. Red shading indicates the timing of optical stimulation (635 nm, 50 ms pulse, 5 Hz, 120 s). Bottom: Ratio (R) calculated by F470/F415 from the upper traces. The baseline ratio (R_0_) is the mean R-value of the pre-stimulation period (-30 s - 0 s). ΔR is the difference between the mean R-value of the late phase during the stimulation period (R_1_, 90 s - 120 s, R_1_) and R_0_. a.u., arbitrary unit. (C, D) Daily rhythms of the baseline ratio (R_0_, open circle) and the ratio during the late phase of stimulation (R_1_, closed circle). Each color represents an individual mouse. C, ChrimsonR; D, mCherry-Control. (E) Comparison of the ΔR/R_0_ %. Optogenetic stimulation of SCN AVP neurons increases VIP-Ca^2+^ during the night in freely moving mice. n = 6 for ChrimsonR, n = 4 for control. ***P < 0.001 by two-tailed Welch’s t-test. Four out of 6 in ChrimsonR and all control animals are from a previously used cohort [12].

### GABA_A_R antagonist increases VIP-Ca^2+^ and inhibits the response of VIP-Ca^2+^ to the optogenetic activation of AVP neurons in SCN slices

To further understand how AVP neurons regulate VIP neurons in the SCN network, we next recorded the VIP-Ca^2+^ response to the optogenetic activation of AVP neurons in coronal slices of the middle SCN along the rostro-caudal axis. ChrimsonR-mCherry and jGCaMP7s were expressed in AVP and VIP neurons of *Avp-Cre; Vip-tTA* mice, respectively, by injecting AAV vectors into the SCN (Fig 5A and 5B). Then, SCN slices were prepared, and an optical fiber was placed close to the ChrimsonR-mCherry-positive region in the SCN for optogenetic activation (Fig 5B).

**Figure 5.**
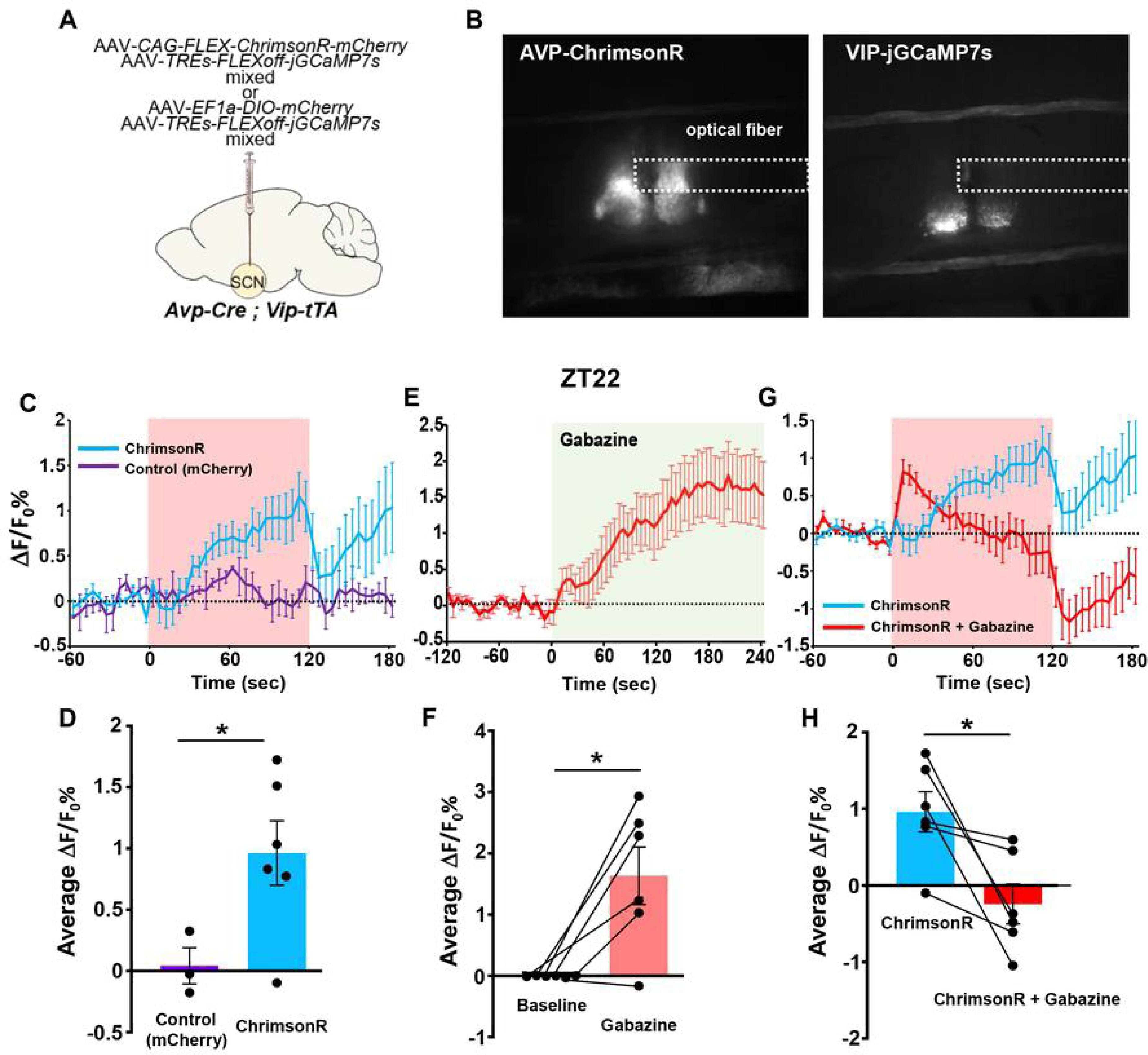
GABA_A_R antagonist increases VIP-Ca^2+^ and inhibits the response of VIP-Ca^2+^ to the optogenetic activation of AVP neurons in slices. (A) Schematic diagram of viral vector injection at the SCN in *Avp-Cre; Vip-tTA* mice for slice recording of VIP-Ca^2+^ and optogenetic stimulation of AVP neurons. (B) A representative coronal section of mice with ChrimsonR expression in AVP neurons (left) and jGCaMP7s expression in VIP neurons (right). White dotted squares indicate the estimated position of the optical fiber. (C) Average traces of the jGCaMP7s signal of SCN VIP neurons with (ChrimsonR, blue) or without (Control, purple) optogenetic stimulation of AVP neurons at ZT22. Red shading indicates the timing of optical stimulation (617 nm, 40 ms pulse, 10 Hz, 120 s). (D) The mean ΔF/F_0_% values during the last 30 s of optogenetic stimulation (90 s – 120 s in C). Values are mean ± SEM. n=3 for control (mCherry) mice, n = 6 for ChrimsonR mice. *P < 0.05 by two-tailed Welch’ s t-test. (E) Average trace of jGCaMP7s signal in VIP neurons after administration of gabazine (GABA_A_R antagonist, 10 μM) via bath perfusion. The green shading indicates when Gabazine was present. (F) The mean ΔF/F_0_% value during the last 60 s of gabazine application (180 s –240 s in E). Baseline is the mean ΔF/F_0_% values 2 min prior to gabazine application (-120 – 0 s). Values are mean ± SEM. n = 6. *P < 0.05 by two-tailed paired t-test. (G) Average traces of the jGCaMP7s signal of SCN VIP neurons upon optogenetic stimulation of AVP neurons at ZT22 without (ChrimsonR, blue) or with (ChrimsonR + Gabazine, red) gabazine application. Red shading indicates the timing of optical stimulation (617 nm, 40 ms pulse, 10 Hz, 120 s). The ChrimsonR traces in (C, blue) and (G, blue) are identical. (H) The mean ΔF/F_0_% values during the last 30 s of optogenetic stimulation (90 s – 120 s in G). Values are mean ± SEM. n = 6. *P < 0.05 by two-tailed paired t-test.

Slices were photo-stimulated around ZT22 when VIP neurons exhibited a strong response in vivo. Optogenetic activation of AVP neurons increased VIP-Ca^2+^, as observed in vivo (Figs 5C, 5D, S4A and S4B). Strikingly, the application of a GABA_A_R antagonist, gabazine, alone raised the baseline VIP-Ca^2^ (Figs 5E, 5F, and S4A), suggesting that VIP neurons are suppressed by GABA during the night. Moreover, the VIP-Ca^2+^ increase induced by the optogenetic activation of AVP neurons was largely inhibited in the presence of gabazine (Figs 5G, 5H, S4A and S4C). These data may suggest that GABA released from AVP neurons indirectly activates VIP neurons by inhibiting intermediate GABA neurons that suppress VIP neurons. On the other hand, the fact that gabazine failed to inhibit the VIP-Ca^2+^ response completely may indicate the presence of parallel pathways mediated by transmitters of AVP neurons other than GABA, such as AVP and other neuropeptides.

### Optogenetic activation of SCN AVP neurons decreases Ca^2+^ in the adjacent non-AVP neurons

If AVP neurons disinhibit VIP neurons, as suggested in the previous section, there should be some population of intermediate GABA neurons that are inhibited by AVP neurons. To test this possibility, we made a spatial map of the Ca^2+^ responses of non-AVP neurons to the optogenetic activation of AVP neurons in SCN slices. To do this, we expressed ChrimsonR-mCherry and jGCaMP7s in AVP and non-AVP neurons, respectively, by injecting AAV-*CAG-FLEX-ChrimsonR-mCherry* and AAV-*EF1-rDIO (reverse DIO)-jGCaMP7s* into the SCN of *Avp-Cre* mice (Fig 6A and 6B). Then, we prepared slices of the middle SCN along the rostro-caudal axis and monitored the jGCaMP7s signal throughout the slices.

**Figure 6.**
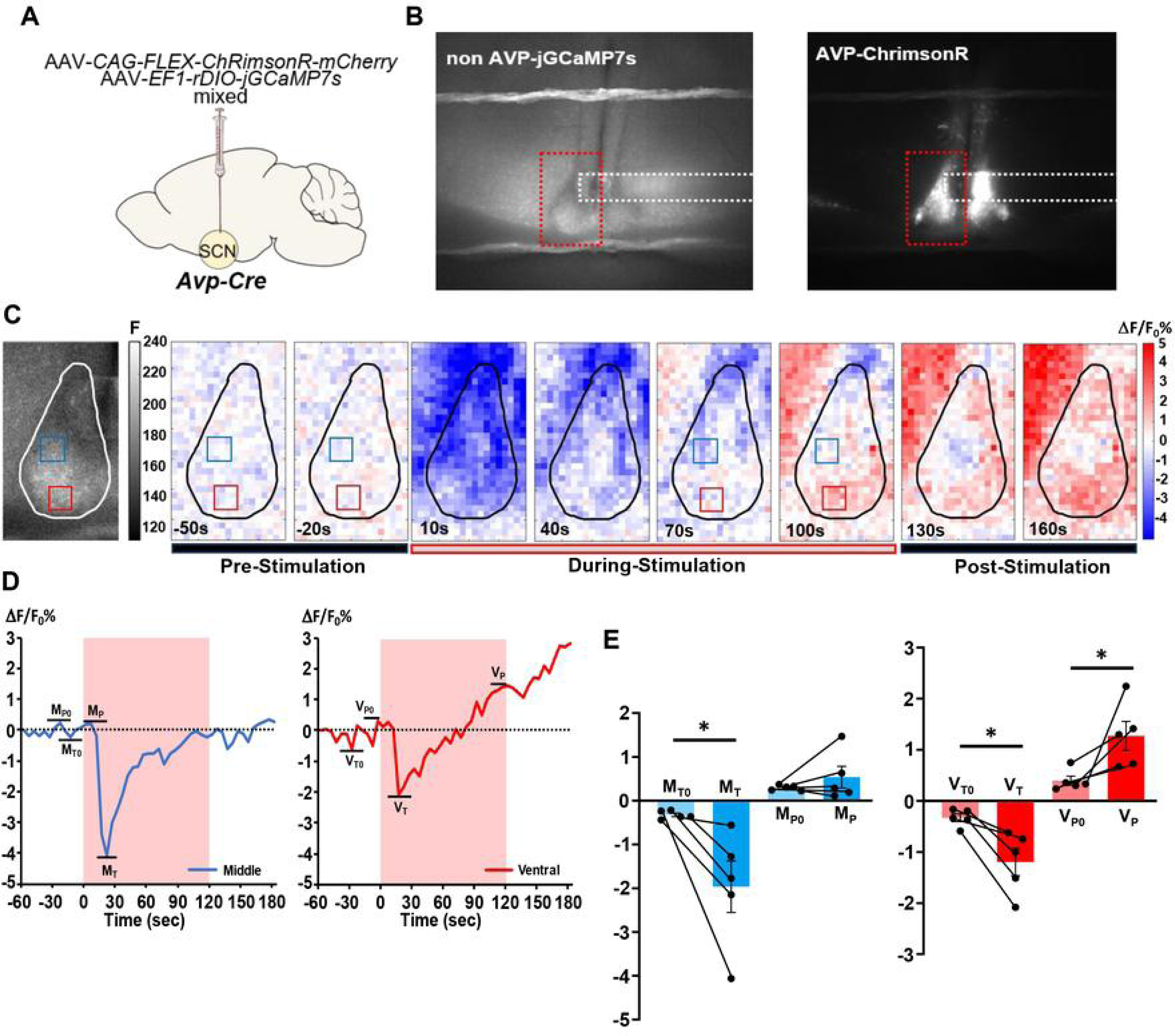
Optogenetic activation of AVP neurons decreases Ca^2+^ in the adjacent non-AVP cells. (A) Schematic diagram of viral vector injection at the SCN in *Avp-Cre* mice for slice recording of SCN non-AVP cellular Ca^2+^ and optogenetic stimulation of AVP neurons. (B) A representative coronal section of mice with jGCaMP7s expression in non-AVP cells (left) and ChrimsonR expression in AVP neurons (right). White dotted squares indicate the estimated position of the optical fiber. The red rectangles in (B) indicate the regions of the enlarged images (C). (C) Left: Representative coronal section of *Avp-Cre* mice with jGCaMP7s expression in non-AVP cells (2.6 μm/pixl). White outline indicates the regions considered as the SCN. Right: Representative pixel-level heat maps (20.8 μm/pixel) showing the jGCaMP7s signal from non-AVP cells in response to the optogenetic stimulation of AVP neurons at ZT22. Optical stimulation was applied from time 0 to 120 s. Blue and red squares on the maps indicate regions (4 x 4 pixels) considered as the middle and ventral regions, respectively, for the subsequent analyses. (D) Representative responses of the jGCaMP7s signals in the middle (left) and ventral (right) regions to the optogenetic activation of AVP neurons. Red shading indicates the timing of optical stimulation (617 nm, 40 ms pulse, 10 Hz, 120 s). M_P0_ and M_T0_ are the highest (peak) and lowest (trough) ΔF/F_0_%-value of the pre-stimulation (-60 s ∼ 0 s) in the middle region, respectively. M_P_ and M_T_ are the highest and lowest ΔF/F_0_%-value during the stimulation (0 s ∼ 120 s). Correspondingly, V_P0_, V_T0_, V_P_, and V_T_ refer to the respective points in the ventral region. (E) Mean responses of the meddle and ventral SCN regions to the optogenetic activation of AVP neurons. Values are mean ± SEM. n = 5. *P < 0.05 by two-tailed Welch’ s t-test.

The jGCaMP7s signals of non-AVP neurons began to decrease shortly (roughly 10 s) after the onset of optogenetic stimulation of AVP neurons in most of the SCN area (Fig 6C). As the stimulation continued, the signals gradually recovered from this inhibitory response and eventually shifted to an increasing response in some areas (Fig 6D). Cells exhibiting such an excitatory response tended to be distributed in the ventral region of SCN, where VIP neurons are located. In contrast, cells in the intermediate area tended to demonstrate larger inhibitory responses with little excitation (Figs 6C-E, S5A and S5B). These results support the idea that AVP neurons indirectly activate VIP neurons via GABA by inhibiting another population of GABA neurons in the SCN network.

## Discussion

In this study, we investigated the GABAergic network through which AVP neurons in the SCN shell regulate VIP neurons in the SCN core to set the timing of locomotor activity. To our knowledge, this is the first mechanistic analysis of the functional regulation from the SCN shell to the core, highlighting the critical role of the GABAergic network within the SCN.

The *Vgat* deficiency in AVP neurons extended the activity time of locomotor activity and compressed the duration of high VIP-Ca^2+^ accordingly to fit in the behavioral rest time. Conversely, the GABA_A_R deficiency in VIP neurons shortened the activity time by reducing morning and evening locomotor activities and lengthened the high VIP-Ca^2+^ duration accordingly. Thus, GABAergic communication from AVP neurons to VIP neurons may account for the inverse correlation between the locomotor activity time and the high VIP-Ca^2+^ duration. Besides, the optogenetic activation of AVP neurons increased VIP-Ca^2+^ both in vivo and ex vivo during the night, which was shown to be inhibited by GABA_A_R antagonism in slices. Furthermore, baseline VIP-Ca^2+^ rose upon GABA_A_R antagonism, and AVP neuronal activation reduced Ca^2+^ in cells located in the SCN area between AVP and VIP neurons. Together with our previous observation that the daily AVP-Ca^2+^ rhythm starts to rise slightly before the onset of VIP-Ca^2+^ [12], these findings raise a model in which GABA released from AVP neurons activates VIP neurons indirectly by suppressing intermediate GABAergic neurons that inhibit VIP neurons, then VIP neurons in turn suppress the locomotor activity to set the rest time, probably indirectly via other neurons that regulate locomotion (S6 Fig). Therefore, the lack of GABA release from AVP neurons may fail to disinhibit VIP neurons in the evening and morning, leading to an earlier onset and later offset of the locomotor activity. Conversely, the lack of GABA_A_R signaling in VIP neurons may reduce their inhibition by the intermediate GABA neurons in the evening and morning, resulting in a later onset and earlier offset of the locomotor activity. It should be noted that not only neurons but also astrocytes may contribute as the intermediate GABA-releasing/regulating cells, as discussed later.

Both the inhibitory and excitatory effects of GABA on SCN neurons have been reported, which may vary with the time of the day, area within the SCN, or photoperiod [14,37–40]. Therefore, we also considered the possibility that GABA from AVP neurons directly excited VIP neurons. However, the fact that gabazine application increased VIP-Ca^2+^ in slices indicated that GABA inhibited most, if not all, VIP neurons. This observation is consistent with a previous study reporting that GABA has a predominantly inhibitory effect on the ventral SCN [37]. In addition, AVP neurons send only sparse projections directly onto VIP neurons [12,34]. On the other hand, there are many GABA neurons that receive abundant AVP fibers and densely project to VIP neurons between AVP-rich and VIP-rich regions in the SCN. Indeed, we confirmed the existence of cells in such intermediate regions that respond to AVP neuronal activation by reducing Ca^2+^. Collectively, the above-mentioned disinhibition pathway via the intermediate neurons is likely the primary regulatory pathway, although an accessory direct GABAergic pathway may exist. In addition, other transmitters, such as AVP, may play an additional role in bridging AVP neurons to VIP neurons.

Whether the presumed intermediate neurons are specialized cells with a specific neurochemical character or a heterogeneous population remains unknown. GRP or calretinin neurons may be good candidates because they are GABA neurons located between AVP and VIP neurons, have contacts of AVP fibers, and make contacts onto VIP neurons [34]. On the other hand, because the SCN is full of many types of GABA neurons, the intermediate neurons may be more broadly distributed spatially and neurochemically. In terms of neuronal activity, at first glance, the presumed intermediate neurons are expected to be more active during the night to inhibit VIP neurons and to be inhibited by AVP neurons. However, MUA recordings have shown that SCN neurons are generally more active during the day, with a peak around midday [1], which apparently contradicts this expectation. Nevertheless, the phase of the firing rhythms of individual SCN neurons is distributed broadly, and indeed, the MUA begins to rise before the (subjective) day and still falls after the onset of the (subjective) night, covering dusk and dawn [25,41,42]. During these transitional time windows, disinhibition of VIP neurons by AVP neuronal GABA may be critical to set the locomotor activity onset and offset precisely. Indeed, the VIP-Ca^2+^ was high at midday even when AVP neurons did not release GABA (*Avp-Vgat^−/−^*), and it was low at midnight even when VIP neurons did not have GABA_A_R (*Vip-GABA_A_R^−/−^*), indicating GABA-independent regulation of daily VIP neuronal activities around their peak and trough, which likely includes the TTFL-driven cellular clocks.

Recent studies using organotypic SCN cultures have nicely demonstrated the critical role of astrocytes in circadian timekeeping by controlling the extracellular GABA levels through rhythmic GABA uptake, GABA release, and glutamate release to stimulate nerve terminal GABA release [19,20,43]. Importantly, astrocytes drive the daily extracellular GABA fluctuation, which is higher at night. Therefore, it is tempting to hypothesize that GABA released from AVP neurons somehow modulates astrocyte activity to reduce GABA levels around VIP neurons.

Although most SCN neurons contain GABA and also express GABA_A_R, they are heterogeneous populations and form assymmetric neural and humoral networks, indicating functional differentiation among cell types. In particular, GABAergic communication between the SCN shell and core is likely to be far from symmetric [37,44,45]. Thus, it is essential to pay attention to the direction of GABAergic signaling under analysis when we investigate the function of GABA in the SCN central clock. To this end, *Vgat* has been deleted to eliminate GABA release specifically in VIP or AVP neurons; the former caused little effect [46,47], while the latter lengthened the activity time of the behavior rhythm [25]. However, until now there has been no means to inhibit GABA reception in a cell type-specific manner without detailed information on the subtype genes expressed in the cells of interest. Here, we achieved a complete cell type-specific blockade of GABA_A_R signaling by deleting all three subtype genes of the requisite β-subunit using in vivo genome editing. A similar strategy to simultaneously disrupt multiple subtype genes would be applicable to study the cell type-specific functions of other ion channels with many subtype members, such as AMPA receptors and potassium channels. Moreover, it is noteworthy that the current strategy of recording the VIP-Ca^2+^ response to optogenetic stimulation of AVP neurons using a combination of Cre/tTA driver mice and Cre/tTA-dependent AAV vectors allowed us to investigate the GABAergic regulatory network in the direction from AVP neurons to VIP neurons.

Although the locomotor activity times were altered, the free-running periods did not change in either *Avp-Vgat^−/−^* or *Vip-GABA_A_R^−/−^* mice. Other reported mice with genetic impairments of VGAT or GABA_A_R also showed free-running periods in the normal range, whereas the amplitudes of the circadian rhythm were reduced to varying degrees [48–50]. We previously showed that clock gene expression rhythms do not alter significantly in either the shell or core of the SCN in *Avp-Vgat^−/−^*mice, suggesting that the TTFLs tick normally in the absence of GABA release of AVP neurons [25]. Thus, AVP neuronal GABA may modify the temporal pattern of VIP neuronal activity with little effect on the TTFL. On the other hand, AVP neurons have been suggested to function as the primary pacesetter cells to determine the SCN ensemble period, which does not require GABA from these neurons [10–12,25]. Therefore, AVP neurons may use GABA to set timers that control the phase relationship of evening and morning locomotor activities on their cellular clocks. Thus, AVP neurons may regulate the pacesetting of the SCN ensemble rhythm and the phase-setting of the evening and morning locomotor activities via partially independent transmitter systems.

## Materials and Methods

### Animals

All experimental procedures were approved by the Kanazawa University Animal Experiment Committee and the Kanazawa University Safety Committee for genetic recombinant experiments. *Avp-Cre* BAC transgenic (C57BL/6J-*Tg(Avp-icre)#Meid*/Rbrc, RBRC12048) [11] and *Vip-tTA* knock-in (B6(Cg)-*Vip^em1(tTA2)Miem^*/Rbrc, RBRC12109) [26] mice were reported previously. *Vip-ires-Cre* (*Vip^tm1(cre)Zjh^/*J, JAX:010908) [33], *Vgat^flox^* (*Slc32a1^tm1Lowl^*/J, JAX:012897) [51], and *Rosa26-LSL-SpCas9-2A-EGFP* mice (B6J.129(B6N)-*Gt(ROSA)26Sor^tm1(CAG-cas9*,-EGFP)Fezh^*/J, JAX:026175) [32] were obtained from Jackson Laboratory. All lines were congenic on C57BL/6. We compared the conditional knockouts with controls whose genetic backgrounds were comparable. *Avp-Cre*, *Vip-ires-Cre*, *Vip-tTA*, and *Rosa26-LSL-SpCas9-2A-EGFP* mice were used in hemizygous or heterozygous condition. We used both male and female mice in our experiment. Mice were maintained under a strict 12-h light/12-h dark cycle in a temperature- and humidity-controlled room and fed ad libitum.

### Viral vector and surgery

The AAV-2 ITR containing plasmids *pGP-AAV-CAG-FLEX-jGCaMP7s-WPRE* (Addgene plasmid #104495, a gift from Dr. Douglas Kim and GENIE Project) [27]. *pAAV*-*TRE-jGCaMP7s* was described previously [52]. In addition, we modified this plasmid to make an improved version by using an EcoRI-HindIII fragment of this plasmid containing *jGCaMP7s* sequence to replace an EcoRI-HindIII fragment containing *ChrimsonR-mCherry* from *pAAV-TRE-ChrimsonR-mCherry* (Addgene #92207, a gift from Alice Ting)[36]. *pAAV-U6-gGabrb1∼3-EF1α-DIO-mCherry*, plasmids for CRISPR-Cas9-mediated *Gabrb1∼3* gene disruption, was generated as follows. The target sites for CRISPR-Cas9 were designed by CRISPOR (http://crispor.tefor.net/) [53]. Two sequences targeting each *Gabrb* gene were selected: *Gabrb1*, 5’-AAGGATATGACATTCGCTTG- 3’ and 5’-CGCATCCCGACGTCCACCGG-3’; *Gabrb2*, 5’-TGACCCTAGTAATATGTCGC-3’ and 5’-ATGTTCATTCCTACGGCCAC-3’; *Gabrb3*, 5’-ATTCGCCTGAGACCCGACTT-3’ and 5’-CGACATCGCCAGCATCGACA-3’.

Oligonucleotides encoding the guide sequences were cloned into the BbsI and BsaI sites of *pX333* (Addgene #64073, a gift from Dr. Andrea Ventura) [54]. Then, a fragment containing two tandem units of *U6-gRNA* was amplified by PCR, using the following primers: 5’-agtacgcgTCGAGCATGCTCGAGAATGG-3’ and 5’-agtacgcgtCGGGTACCCCATTTGTCTGC-3’, and cloned into the MluI site of *pAAV-EF1a-DIO-mCherry* (a gift from Dr. Bryan Roth) as described previously [52]. *pAAV-U6-gGabrb2-EF1α-DIO-mCherry* happened to contain two copies of the amplified fragment. *pAAV-U6-gControl-EF1α-DIO-mCherry* contains spacer sequences from *pX333* (Addgene plasmid #64073, a gift from Dr. Andrea Ventura) instead of gRNA sequences for *Gabrb3* genes. *pAAV*-*CAG-FLEX-ChrimsonR-mCherry* was described previously (9). As a negative control for the optogenetic study, we injected AAV-*EF1α-DIO-mCherry* or AAV-*EF1α-DIO-hM3Dq-mCherry*, which was generated with plasmids *pAAV-EF1α-DIO-mCherry* or *pAAV-EF1α-DIO-hM3Dq-mCherry* provided by Dr. Bryan Roth, University of North Carolina [55]. *pAAV*-*TRE-FLEXoff-jGCaMP7s* was made by replacing an EcoRI-SpeI fragment containing *ChrimsonR-mCherry* of *pAAV-TRE-ChrimsonR-mCherry* with an EcoRI-XbaI fragment containing *jGCaMP7s* from *pGP-AAV-CAG-FLEX-jGCaMP7s-WPRE*. *pAAV*-*EF1-rDIO-jGCaMP7s* was made by replacing an AscI-NheI fragment containing *ChR2-EYFP* of *pAAV-DIO-hChR2(H134R)-EYFP-WPRE-pA* (provided by Dr. Karl Deisseroth, Stanford University) with a *jGCaMP7s* cDNA fragment amplified by PCR from *pGP-AAV-CAG-FLEX-jGCaMP7s-WPRE,* using the following primers: 5’-ataggcgcGCCACCATGGGTTCTCATCA-3’ and 5’-gcgactagTCACTTCGCTGTCATCATTTG-3’. Note that gene expression from AAV-*TRE-FLEXoff-jGCaMP7s* and *AAV*-*EF1-rDIO-jGCaMP7s* stops upon Cre-mediated recombination. Recombinant AAV vectors (AAV2-rh10) were produced using a triple-transfection, helper-free method and purified as described previously [11]. The titers of recombinant AAV vectors were determined by quantitative PCR: AAV-*CAG-DIO-jGCaMP7s*, 3.4 × 10^13^; AAV-*TRE-jGCaMP7s*, 6.3 × 10^11^; AAV-*TRE-jGCaMP7s* (improved), 5.8 × 10^12^; AAV-*TRE-FLEXoff-jGCaMP7s*, 1.48 × 10^13^; AAV-*EF1-rDIO-jGCaMP7s*, 1.5 × 10^13^; AAV*-U6-gGabrb1-EF1α-DIO-mCherry*, 6.06 × 10^12^; AAV*-U6-gGabrb2-EF1α-DIO-mCherry*, 7.42 × 10^12^; AAV*-U6-gGabrb3-EF1α-DIO-mCherry*, 6.56 × 10^12^; AAV-*U6-gControl-EF1α-DIO-mCherry*, 2.6 × 10^12^; AAV-*EF1α-DIO-hM3Dq-mCherry*, 4.5 × 10^12^; AAV-*CAG-FLEX-ChrimsonR-mCherry*, 1.5 × 10^13^; and AAV-*EF1α-DIO-mCherry*, 5.2 × 10^12^ genome copies/ml. Stereotaxic injection of AAV vectors was performed as described previously [11]. Two weeks after surgery, we began monitoring the mice for their locomotor activity.

### In vivo fiber photometry

We used 8 *Avp-Vgat^−/−^* × *Vip-tTA* (*Avp-Cre; Vgat ^flox/flox^; Vip^wt/tTA^*) mice, 5 control mice (*Avp-Cre; Vgat^wt/flox^; Vip^wt/tTA^*) and 12 *Vip-ires-Cre; Rosa26-LSL-SpCas9-2A-EGFP* mice for the *GABA_A_R* disruption study (5 or 7 for *gGabrb1-3* or *gControl*). The mice were anesthetized by administering a cocktail of medetomidine (0.3 mg/kg), midazolam (4 mg/kg), and butorphanol (5 mg/kg) and were secured at the stereotaxic apparatus (Muromachi Kikai). Lidocaine (1%) was applied for local anesthesia before making the surgical incision. We drilled small hole in the exposed region of the skull using a dental drill. We injected 0.5 to 1.0 μL of the virus (AAV*-U6-gGabrb1-3-EF1α-DIO-mCherry ; AAV-CAG-FLEX-jGCaMP7s* mixed *(gGabrb1: gGabrb2: gGabrb3: jGCaMP7s=1:1:1:2)* or AAV*-U6-gControl-EF1α-DIO-mCherry ; AAV-CAG-FLEX-jGCaMP7s* mixed or *AAV-TRE-jGCaMP7s*) (flow rate = 0.1 μL/min) at the right or bilateral of SCN (posterior: 0.5 mm, lateral: 0.25 mm, depth: 5.7 mm from the bregma) with a 33 G Hamilton Syringe (1701RN Neuros Syringe, Hamilton) to label VIP neurons. Subsequently, we placed an implantable optical fiber (400 μm core, N.A. 0.39, 6 mm, ferrule 2.5 mm, FT400EMT-CANNULA, Thorlabs or RWD) above the SCN (posterior: 0.2 mm, lateral: 0.2 mm, depth: 5.2 to 5.4 mm from the bregma) with self-adhesive resin cement (Super-bond C&B, Sun Medical or RelyX^TM^ Unicem2 Automix, 3M ESPE). The cement was painted black. Atipamezole (0.3 mg/kg) was administered postoperatively to reduce the anesthetized period. The mice were used for experiments 2 to 8 weeks after the virus injection and optical fiber implantation. Their ages ranged from 3 to 14 months old, including both males and females.

A single-color fiber photometry system (COME2-FTR, Lucir) was used to record the Ca^2+^ signal of SCN neurons in freely moving *Avp-Vgat^−/−^* mice (Figs 1 and S1) [12,56]. Fiber-coupled LED (M470F3, Thorlabs) with LED Driver (LEDD1B, Thorlabs) was used as an excitation blue light source. The light was reflected by a dichroic mirror (495 nm), went through an excitation bandpass filter (472/30 nm), and then to the animal via a custom-made patch cord (400 um core, N.A. 0.39, ferrule 2.5 mm, length 50 cm, COME2-FTR/MF-F400, Lucir) and the implanted optical fiber. We detected the jGCaMP7s fluorescence signal by a photomultiplier through the same optical fibers and an emission bandpass filter (520/36 nm); furthermore, we recorded the signal using Power Lab (AD Instruments) with Lab Chart 8 software (AD Instruments). The excitation blue light intensity was 10 to 30 μW at the tip of the patch cord of the animal side. We recorded the same for 30 s every 10 min for 2 weeks to reduce photobleaching. During the recording, the mouse was housed in a 12-h light-dark cycle for more than 5 days (LD condition) and then moved to continuous darkness for approximately 10 days (DD condition) in a custom-made acrylic cage surrounded by a sound-attenuating chamber. A rotary joint for the patch cord was stopped during the recording to prevent artificial baseline fluctuation. The animal’s locomotor activity was monitored using an infrared sensor (Supermex PAT.P and CompACT AMS Ver. 3, Muromachi Kikai).

The detected GCaMP signal was averaged within a 30-s session [25]. To detrend the gradual decrease of the signal during recording days, ±12 h average from the time (145 points) was calculated as baseline (F). The data were subsequently detrended by the subtraction of F (ΔF). Then, the ΔF/F value was calculated. To determine VIP-Ca^2+^ onset and VIP-Ca^2+^ offset, ΔF/F were smoothened with a 21-point moving average, then the local maximum and minimum time points within one day were determined. Then, the time points crossing the 50% amplitude (ΔF/F maximum – minimum value) were defined as VIP-Ca^2+^ onset and offset, respectively. The middle of the time points between the VIP-Ca^2+^ onset and offset were defined as the VIP-Ca^2+^ midpoint. Additionally, the intervals between VIP-Ca^2+^ onset and offset were defined as high Ca^2+^ duration. A double-plotted actogram of jGCaMP7s signal was designed by converting all ΔF to positive values by subtracting the minimum value of ΔF. Subsequently, these values were multiplied by 100 or 1,000 and rounded off. The plots were made via ClockLab (Actimetrics) with normalization in each row.

Another dual-color fiber photometry system (FP3002, Neurophotometrics) was used to record the calcium signal of SCN neurons in freely moving *Vip-GABA_A_R^−/−^* mice (Figure 3) [12,57,58]. Excitation light sources were a 470-nm LED for detecting calcium-dependent jGCaMP7s fluorescence signal (F470) and a 415-nm LED for calcium-independent isosbestic fluorescence signal (F415). The duration of excitation lights is 50 ms, and the onsets of the excitation timing of LEDs were interleaved. The lights passed through excitation bandpass filters, dichroic mirrors, and then to the animal via fiber-optic patch cords (BBP(4)_400/440/900-0.37_1m_FCM-4xFCM_LAF, MFP_400/440/LWMJ-0.37_1m_FCM-ZF2.5_LAF, Doric Lenses) and the implanted optical fiber. Subsequently, both signals were detected using a CMOS camera through the optical fibers, dichroic mirrors, and emission bandpass filters. The recorded signals were acquired using Bonsai software, with a sampling rate of 10 Hz for each color. The excitation intensities of the 470-nm and 415-nm LED at the animal side’s patch cord tip were from 60 μW to 110 μW. We recorded the same for 30 s every 10 min for 3 weeks to reduce photobleaching.

The detected GCaMP signal was averaged within a 30-s session. Ratio (R) was defined as the ratio between F470 and F415 (F470/F415) for calibration and reducing motion artifacts. To detrend the gradual decrease of the signal during recording days, ±12 h average from the time (145 points) was calculated as baseline (R_0_). The data were subsequently detrended by the subtraction of R_0_ (ΔR). Then, the ΔR/ R_0_ value was calculated. To determine VIP-Ca^2+^ onset and VIP-Ca^2+^ offset, ΔR/R_0_ were smoothened with a 21-point moving average, then the local maximum and minimum time points within one day were determined. Then, the time points crossing the 50% amplitude (ΔR/R_0_ maximum – minimum value) were defined as VIP-Ca^2+^ onset and offset, respectively. The middle of the time points between the VIP-Ca^2+^ onset and offset were defined as the VIP-Ca^2+^ midpoint. Additionally, the intervals between midpont phases were defined as the periods [11] and the intervals between VIP-Ca^2+^ onset and offset were defined as high Ca^2+^ duration. A double-plotted actogram of GCaMP signal was designed by converting all R values. The plots were made via ClockLab (Actimetrics).

Ca^2+^ activity profile analyses were performed via ClockLab (Actimetrics). Before analyzing with the ClockLab, all R or ΔF values were normalized by subtracting the minimum value of R or ΔF, dividing the result by the difference between the maximum and minimum R or ΔF values, and then multiplying by 100 or 1000. Subsequently, the last 5 days in the LD condition and last 5 or 10 days in the DD condition were selected using ClockLab for Ca^2+^ activity profile analysis. The resulting profiles were further normalized in the same way to generate the final Ca^2+^ activity profile. CT12 was determined as the onset of locomotor activity. VIP-Ca^2+^ CT0 was determined as the onset of VIP-Ca^2+^ activity. High Ca^2+^ duration (20% amplitude) was the interval between VIP-Ca^2+^ onset and offset defined as the time points crossing the 20% amplitude of normalized calcium activity profile with VIP-Calcium circadian time, rather than 50% as described above.

During the fiber photometry recordings, the animal’s locomotor activity was monitored using an infrared sensor (Supermex PAT.P and CompACT AMS Ver. 3, Muromachi Kikai) in 1-min bins, then 10-min bins were made by analysis. A double-plotted actogram of locomotor activity was also prepared and overlaid on that of the GCaMP. The onset and offset of locomotor activity were determined using the actogram of locomotor activity. Initially, we attempted to automatically detect the onset and offset; however, it was followed by a manual visual inspection and modifications by the experimenter. The intervals between locomotor activity onset and offset were defined as locomotor activity time.

We confirmed the jGCaMP7s or gRNA (mCherry) expression and the position of the optical fiber by slicing the brains into 30 or 100 μm coronal sections using a cryostat (Leica). The sections were mounted on glass slides with a mounting medium (VECTASHIELD HardSet with DAPI, H-1500, Vector Laboratories or Dako Fluorescence Mounting Medium, Agilent Technologies) and observed via epifluorescence microscope (KEYENCE, BZ-9000E).

### Behavioral analyses

Male and female *Vip-GABA_A_R^−/−^* (*Vip-ires-Cre; Rosa26-LSL-SpCas9-2A-EGFP* injected bilaterally with AAV*-U6-gGabrb1∼3-EF1α-DIO-mCherry* mixture) and control (injected with control AAV) (Figure 2C) mice, aged 8 to 20 weeks, were housed individually in a cage placed in a light-tight chamber (light intensity was approximately 100 lux). Spontaneous locomotor activity (home-cage activity) was monitored by infrared motion sensors (O’Hara) in 1-min bins as described previously [12]. Actogram, activity profile, and χ^2^ periodogram analyses were performed via ClockLab (Actimetrics). The free-running period and amplitude (Qp values) were calculated for the last 10 days in constant darkness (DD) by periodogram. The onset of locomotor activity, defined as CT12, was calculated from the daily activity profile of the same 10 days in DD using the median activity level as a threshold for onset detection (Fig 2D and 2E). The onset and offset of locomotor activity used to calculate the locomotor activity time (Fig 2F) were determined using the actogram. Initially, we attempted to automatically detect the onset and offset; however, it was followed by a manual visual inspection, along with modifications by the experimenter.

### In vivo optogenetic stimulation with fiber photometry

We used 10 *Avp-Cre; Vip-tTA* mice (*n =* 6 for ChrimsonR, *n* = 4 for control, both male and female). Some animals in this section are from a previously used cohort [12]. We injected 1.0 μL of the mixture of viruses (AAV-*TRE-jGCaMP7s* with AAV-*CAG*-*Flex-ChrimsonR-mCherry* or AAV-*EF1α*-*DIO-hM3Dq-mCherry* and AAV-*TRE-jGCaMP7s*) into the right SCN (posterior: 0.5 mm, lateral: 0.25 mm, depth: 5.7 mm from the bregma) and then implanted an optical fiber (400 μm core, N.A. 0.39, 6 mm, Thorlabs) above the SCN (posterior: 0.2 mm, lateral: 0.2 mm, depth: 5.3 mm from the bregma) with dental cement. The mice were used for experiments more than 2 weeks after the surgery.

The dual-color fiber photometry system (FP3002, Neurophotometrics) was used to record the Ca^2+^ signal of SCN neurons with optogenetic stimulation in freely moving mice, as described above in “In vivo fiber photometry” (Fig 4) [57,58]. The excitation intensities of the 470-nm and 415-nm LEDs at the tip of patch cord on the animal side ranged 130 μW and 90μW, respectively. Additionally, a 635-nm red laser (inside the fiber photometry system FP3002) was transmitted through the same optical fibers with an intensity of 2 mW. During the 720-s recording every 2 h, optical stimulation (635 nm, 50 ms pulse, 5 Hz, 120 s, 600 pulses) was applied in the middle of the recording. Throughout the experiment, the mice were housed in a custom-made acrylic cage surrounded by a sound-attenuating chamber and maintained in a 12-h LD cycle.

The recorded data were interleaved to eliminate artifacts caused by red laser stimulation, and half of it was discarded. Consequently, the final sampling rate for the jGCaMP7s fluorescence signals at the 470-nm light excitation (F470) and the 415-nm excitation (F415) was 5 Hz. Ratio (R) was defined as the ratio between F470 and F415 (F470/F415) for calibration and reducing motion artifacts. The baseline ratio (R0) is the mean R-value of the pre-stimulation period (-30 s - 0 s). ΔR is the difference between the mean R-value of the late phase during the stimulation period (90 s - 120 s, R1) and R0. After the recordings were completed, we confirmed the jGCaMP7s and ChrimsonR-mCherry expressions and the position of the optical fiber by histology.

### Slice electrophysiology

*Vip-GABA_A_R^-/-^* (made as described above) mice were compared to control (*Vip-ires-Cre; Rosa26-LSL-Cas9-2A-EGFP* with AAV-EF1a-DIO-mCherry injected, 1.0 μL bilateral of SCN) mice. Both male and female mice aged 8 to 20 weeks were used. Coronal brain slices (250 µm thick), including the SCN, were prepared with a linear-slicer (NLS-MT, Dosaka EM), as described previously [59]. Under an upright fluorescence microscope (Olympus, BX51WI), we visually identified VIP neurons with EGFP fluorescence in the ventral SCN and non-VIP neurons without the fluorescence in the dorsal SCN. The gRNA AAV-infected VIP neurons were identified by additional mCherry fluorescence. For recording of non-glutamatergic spontaneous postsynaptic currents (PSCs), the slices were continuously perfused with an artificial CSF (ACSF) with the following composition (in mM): 125 NaCl, 2.5 KCl, 2 CaCl2, 1 MgSO4, 26 NaHCO3, 1.25 NaH2PO4, 10 d-glucose, equilibrated with 95% O2 and 5% CO2, kept at 31 ± 1 °C, and further mixed with 10 μM 6-cyano-7-nitroquinoxaline-2,3 dione disodium (CNQX) and 25 μM D-(-)-2-amino-5-phosphonopentanoic acid (D-AP5) to block fast glutamatergic transmission. Whole-cell voltage-clamp recordings were made at −60 mV with borosilicate glass electrodes (5–6 MΩ) filled with an internal solution containing the following components (in mM): 87 CsCl, 20 TEA-Cl 10 HEPES, 10 EGTA, 0.5 CaCl2, 4 MgCl2, 2 QX314, 4 Na-ATP, 0.4 Na-GTP, and 10 phosphocreatine (pH 7.3, adjusted with CsOH). A combination of an EPC10/2 amplifier and Patchmaster software (HEKA) was used to control membrane voltage and data acquisition. Series resistance was compensated routinely by 80%. The non-glutamatergic PSCs completely disappeared in the presence of 10 μM gabazine (SR95531, Tocris), showing that they were mediated by GABA_A_ receptors [25]. The GABAergic PSCs recorded for 1-3 min were analyzed using the MiniAnalysis program (Synaptosoft). The events were picked by an amplitude threshold of 5 pA and confirmed visually to have a typical PSC waveform.

### Ex vivo optogenetic stimulation and calcium imaging

We used 9 *Avp-Cre; Vip-tTA* mice (*n =* 6 for ChrimsonR, *n* = 3 for control) for VIP-Ca^2+^ recording and 5 *Avp-Cre* mice for recording Ca^2+^ in non-AVP neurons (aged 8 to 20 weeks, including both males and females). We injected 0.6 μL of the mixture of viruses (VIP-Ca^2+^: AAV-*TRE-FLEXoff-jGCaMP7s* with AAV-*CAG*-*Flex-ChrimsonR-mCherry* or AAV-*EF1α*-*DIO-mCherry,* non-AVP-Ca^2+^: AAV-*EF1α-rDIO-jGCaMP7s* with AAV-*CAG-Flex-ChrimsonR-mCherry*) into the SCN bilateral (posterior: 0.5 mm, lateral: 0.25 mm, depth: 5.7 mm from the bregma). Coronal brain slices (300 µm thick), including the SCN, were prepared as described in “Slice electrophygiology.” For this experiment, mice were sacrificed at ZT19–20 in darkness under a red light. The slices were placed in an imaging chamber and continuously perfused with the ACSF. Before the recordings started around projected ZT22, the slices were pre-incubated in the experimental environment for at least one hour, which was critical to observe VIP-Ca^2+^ response to the optogenetic stimulation. Under the upright fluorescence microscope (Olympus, BX51WI), we visually identified AVP neurons with ChrimsonR-mCherry or mCherry (control) fluorescence in the dorsomedial SCN and VIP neurons with jGCaMP7s fluorescence in the ventral SCN. We imaged jGCaMP7s fluorescence every 5 sec through an optical filter set (an excitation filter 470-495 nm, a dichroic mirror 505 nm and an emission filter 510- 550 nm) and a digital CMOS camera (Prime BSI Express, Teledyne Vision Solutions) with MetaFluor software.

After the GCaMP signal had stabilized, we optogenetically stimulated SCN AVP neurons for 2 min (617 nm, 10 Hz, 40 ms duration) through an optic fiber (200 μm core, N.A. 0.39, 6 mm, ferrule 1.25 mm, FT200EMT-CANNULA; Thorlabs) positioned near the AVP neurons and connected to a high-powered LED under the control of Digital Stimulator (WPI DS8000B) and LED driver (THORLABS). Gabazine (10 μM) was applied to ACSF to verify GABA_A_R involvement. For VIP-Ca^2+^, we measured the fluorescence values of the entire jGCaMP7s-expressing ROIs using MetaFluor software, detrended the traces by division, and calculated the change in fluorescence over baseline fluorescence (ΔF/F_0_%) as (Fi-F_0_) * 100 / F_0_. The average F values 1 or 2 min before the optogenetic stimulation or blockers perfusion were defined as the baseline (F_0_).

For non-AVP-Ca^2+^, we selected unilateral SCN regions (150 x 260 pixels, 2.6 μm/pixel) for further analysis. Images were analyzed using MATLAB. ΔF/F_0_% values of individual pixels were calculated as described above. The resolution of images was then adjusted to 20.8 μm/pixel, and square regions of interest (ROIs, 4 × 4 pixels) were defined in the middle and ventral of the SCN. M_P0_ and M_T0_ are the highest and lowest ΔF/F_0_% values, respectively, of the pre-stimulation period (-60 s ∼ 0 s) in the middle region. M_P_ and M_T_ are the highest and lowest ΔF/F_0_% values during the stimulation period (0 s ∼ 120 s). Correspondingly, V_P0_, V_T0_, V_P_, and V_T_ refer to the respective points in the ventral region.

### Statistical analysis

All results are expressed as mean ± SEM. For comparisons of two groups, two-tailed Student’s t test or Welch’s *t* test was performed. For comparisons of multiple groups with no difference of variance, two-way repeated measures ANOVA followed by post hoc two-tailed Student’s t-test were performed. For comparisons of multiple groups with difference of variance, nonparametric tests, Kruskal–Wallis test with post hoc Dunn’s test were performed. All *P* values less than 0.05 were considered as statistically significant. Only relevant information from the statistical analysis was indicated in the text and figures.

## Acknowledgements

We thank H. Okamoto for the *Avp-Cre* mouse; S. Horike and T. Daikoku for the *Vip-tTA* mouse; Z. J. Huang for the *Vip-ires-Cre* mouse; F. Zhang for the *Rosa26-LSL-SpCas9-2A-EGFP* mouse; B.B. Lowell for the *Vgat^flox^* mouse; Penn Vector Core for *pAAV2-rh10*; D. Kim & GENIE Project for *pGP-AAV-CAG-FLEX-jGCaMP7s-WPRE*; A. Ting for *pAAV-TRE-ChrimsonR-mCherry*; A. Ventura for *pX333*; B. Roth for *pAAV-EF1a-DIO-mCherry* and *pAAV-EF1α-DIO-hM3Dq-mCherry*; and *K. Deisseroth* for *pAAV-DIO-hChR2(H134R)-EYFP-WPRE-pA*; We thank all lab members, including M. Kawabata and Y. Nishiwaki. This work was supported in part by JSPS KAKENHI Grant Numbers JP24KJ1189 (Y.P.); JP23K06345 (Y.T.); JP22K20738 (A.M.); JP24K02137 (T.M.); JP23K24064; JP25K02440; the Takeda Science Foundation; the Terumo Life Science Foundation; the Research Foundation for Opto-Science and Technology; the Koyanagi Foundation (M.M.); and JST SPRING Grant Number JPMJSP2135 (M.W., Y.P.).

## Author contributions

Y.P., Y.T. and M.M. designed research; Y.P., Y.T., T.M., M.W., and A.M. performed research; Y.P., Y.T., T.M., and M.W. analyzed data; Y.P., Y.T., and M.M. wrote the paper.

## Supporting Information

S1 Fig. Circadian rhythms of the behavior and VIP-Ca^2+^ in *Avp-Vgat^−/−^* mice

S2 Fig. Disruption of GABA_A_R in SCN VIP neurons by in vivo genome editing

S3 Fig. Circadian rhythms of the behavior and VIP-Ca^2+^ in *Vip-GABA_A_R^−/−^* mice used for fiber photometry recordings

S4 Fig. Response of VIP-Ca^2+^ to the optogenetic activation of AVP neurons in SCN slices

S5 Fig. Response of non-AVP cells to the optogenetic activation of AVP neurons in coronal SCN slices

S6 Fig. A model of how the GABAergic network from AVP neurons to VIP neurons in the SCN sets the activity/rest time of the circadian behavior rhythm

